# STIX: Long-reads based Accurate Structural Variation Annotation at Population Scale

**DOI:** 10.1101/2024.09.30.615931

**Authors:** Xinchang Zheng, Murad Chowdhury, Behzod Mirpochoev, Aaron Clauset, Ryan M Layer, Fritz J Sedlazeck

## Abstract

The prioritization of Structural Variants (SV), which is needed to rank and identify potential pathogenic alleles, is still in its infancy. This is exemplified over gnomAD only being able to annotate 33.5% of GIAB SVs. To overcome this, we present the first long-read based annotation resource for both GRCh38 and CHM13-T2T reference using STIX. In contrast to previous methods, STIX indexes SV-informative long-reads themselves, can thus be easily extended and accurately annotate all SV types including insertions. STIX successfully annotated 95.9% of GIAB Tier1 SVs. STIX further improved cancer based SV prioritization by highlighting 3,563 SV from COSMIC being common in the population. We further showcase that mosaic SV can be independently gained and may be widely spread throughout the population. This highlights the need for accurate SV population frequency annotation to further facilitate the adoption of SV via long-read sequencing in medical research and clinical applications.

## Introduction

Structural variants (SVs) are genomic alterations larger than 50 bp, typically including deletions, duplications, insertions, inversions, translocations, and their combinations^1,2^. SVs impact mendelian, neurological, and cardiovascular diseases, ^3–5^, and are the driver mutations in many cancers including leukemia^6^, lung cancer^7^ and triple-negative breast cancer^8^. SVs often occur in the cancer genomes at lower frequencies but dynamically evolve under selection pressure, resulting in highly heterogeneous tumor SV profiles^8–11^. SV detection is improving, with clear benefits from using long-read sequencing^2,12–15^. However, previously published genomic annotation tools such as VEP^16^, ANNOVAR^17^, or SnpEff^18^ mainly focus on small variants, while SV prioritization and interpretation are lacking behind, which limits our ability to determine the role of SVs in disease studies.

Population frequency annotation is an effective approach to prioritize SVs since a variant that is common in the population is less likely to be pathogenic^19^. Current approaches such as dbVar^20^ and gnomAD^21,22^ report SV population frequencies, but are limited for two main reasons. First, they are based on short-read sequencing, which includes only a fraction of the SVs when compared to using long-reads^12^. Second, they are built upon merged and heavily filtered VCFs that do not depict a harmonized representation^23^. Merging based on the rough overlap (e.g., 70%)^24^ can add significant bias when the caller and sequencing technologies (e.g insert size) vary. Over-merging can occur due to imprecise thresholds, which leads to a wrong interpretation of the allele itself^25^. While the identification of SVs is constantly improving, the population frequency annotation remains challenging, resulting in ineffective prioritization and impact prediction^26^.

Long-read sequencing has significantly improved our understanding of the occurrence and impact of SVs^1,27^. It has better alignments in repetitive regions such as tandem repeats where 70% or more of SV are located^1,28^; is able to span the alleles; and has generally less sequencing biases than short read sequencing^1^. Despite the increased sequencing costs and sample requirements^1^, several current efforts in population long-read sequencing have emerged to provide novel SV insights^2,29–31^, and the time is right to generate a harmonized SV annotation resource to improve SV population frequency estimation. The challenge is the harmonization and the future readiness of such a resource as current annotation methods still require version specific callers and other filtering heuristics. To overcome some of these challenges, we recently introduced STIX for short-read sequencing that overcomes limitations of merged VCF files, by indexing and searching SV informative reads directly^26^. While the initial version of STIX improved SV prioritization, it did not support insertions nor was it able to utilize long-read sequencing data.

In this study, we expanded STIX to long-reads, allowing for the annotation of SV with population-scale evidence across different long-read sequencing platforms. In this update, we fully support all types of SVs, including insertions, and benchmarked the performance of STIX not only for germline SV annotation but also demonstrated that STIX can also improve mosaic SV annotation. To accomplish this, we generated the first long-read annotation resource across both GRCh38 and CHM13-T2T^32^ reference utilizing 1,108 samples spanning all ethnicities. We demonstrate the utility of this catalog together with STIX for variant prioritization across cancer and somatic variant calling experiments. STIX has now solved the long-read annotation issues and enables a future ready resource for the genomic community that can be easily accessed, extended and updated.

## Results

### Comprehensive and accurate SV annotation using STIX

SV annotation is important for rare disease research, medical applications, and genotype-phenotype related studies^33–35^. Our extensions of STIX to long-reads builds on our existing short-read platform^26^ by supporting all SV types. **Fig. 1A** summarizes the three key steps of STIX. In the first step, STIX screens a bam file using a new implementation of excord called excord-lr (**see methods**). Excord-lr scans the bam file’s CIGAR string and split-read signal to find potential SV information. The results are then stored in a bed file for each sample, reducing the storage burden by 99.85%. We have expanded excord-lr to include support for insertions, which are encoded in long-reads in three different ways based on the size of the event (**Fig. 1B**). Small insertions are included in the CIGAR string, split read events represent mid-sized insertions, and insertions larger than the read length generate a single primary alignment with large unaligned segments.

**Fig. 1.**
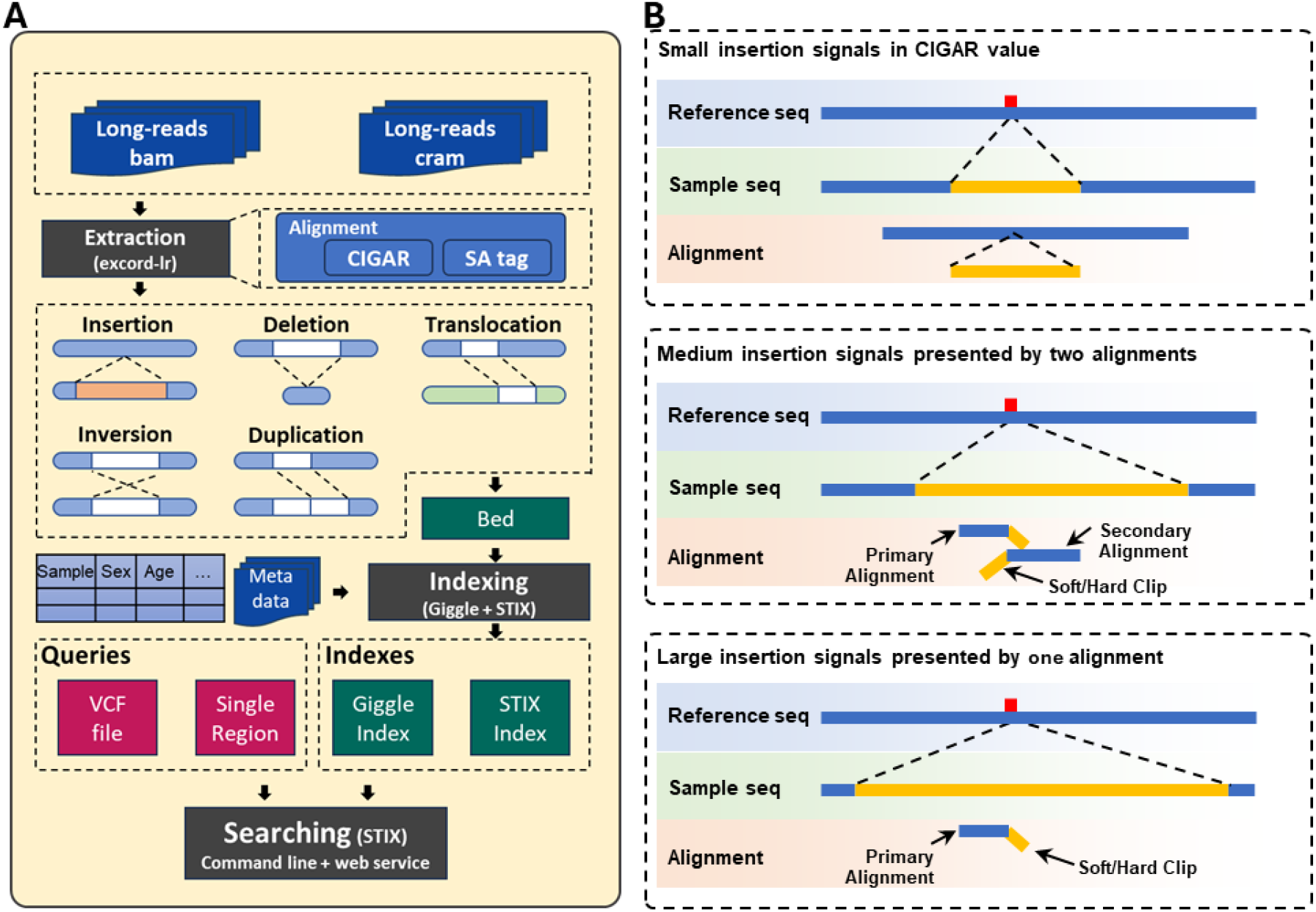
Overview of STIX: **A)** An overview of the main steps in STIX SV annotation **B)** Insertion signals extracted by excord-lr in long-reads dataset are classified by size. Small insertions are in CIGAR string (top). Longer insertions are either one primary alignment and one supplementary alignment (middle), or a single primary alignment (bottom).

In the second step, STIX creates a compressed index^36^ of the previously extracted SV data. Sample-level metadata, such as ethnicity, gender, and phenotype can be added to the index and later retrieved. To enable the resource’s future-ready growth, we have implemented a sharding approach that allows the simultaneous handling of independent indexes and queries.

The third step annotates novel SVs by querying the previously generated indexes. Here STIX assesses if a SV is in the indexed population based on SV type, position, and length. In this step, we can integrate over short- and long-reads and annotate all SV types, including insertions. See methods for a detailed description about the individual steps. STIX is open source and available at https://github.com/ryanlayer/stix together with the annotation index at https://stix.colorado.edu.

### Performance assessment of germline structural variation annotation

STIX returns the population evidence depth for a query SV. To account for the imprecision in SV calling, STIX uses a *padding* parameter to determine how many bases up and downstream of a variant to consider. To derive this threshold and the minimum evidence depth for reporting an SV, we used the HG002 SV benchmark from the Genome in a Bottle (GIAB) project (Tier1GIAB SVs) and created an HG002 STIX index for long-read sequencing data from Oxford Nanopore (ONT) and PacBio HiFi (Pb-HiFi)^37^. SV calls in the benchmark and recovered by STIX were considered true positives (TP). Calls not recovered by STIX are false negatives (FN). Using the GIAB phased assembly SV pipeline, we generated calls for HG00733^28,25^. Calls that were specific to HG00733 and recovered by STIX in the HG002 indexes were considered false positives (FP) (**see methods**).

We tested the robustness of STIX using SV benchmarks from three reference genomes, including the well established HG002 (v0.6) on GRCh37 and two prototype SV benchmarks from GIAB for GRCh38 and CHM13-T2T (V0.012-20231107). For FP, we called SVs in HG00733 based on its assembly compared to GRCh37, GRCh38 and CHM13-T2T, excluded common SV with HG002 and queried these HG00733 private SV against the HG002 index.

As expected, TP and FP rates decreased as the minimum number of supporting reads and the padding increased(**Fig. 2A**). Interestingly, the padding parameter impacted TP, but had minimal effect on FP. We observed slightly different impacts on parameter choice across the two sequencing platforms(**Supplementary FIgure S5**). Pb-HiFi exhibited consistent performance independent of the padding, while ONT benefited from larger paddings. This difference may be due to higher error rates given that the GIAB ONT SVs were produced with the older R9 basecaller with higher error rate^38,39^. Based on these results, we recommend a padding of 100 bp and a minimum read support of 5 reads for germline variants. With these parameters, STIX achieved an overall TP rate of 81.82% and a FP rate of 3.44%. These parameters further minimized the differences between Pb-HiFi and ONT across all three reference genomes(**Fig. 2E**, additional comparisons in **Supplementary Section 1** and **Supplementary Table S2**).

**Fig. 2.**
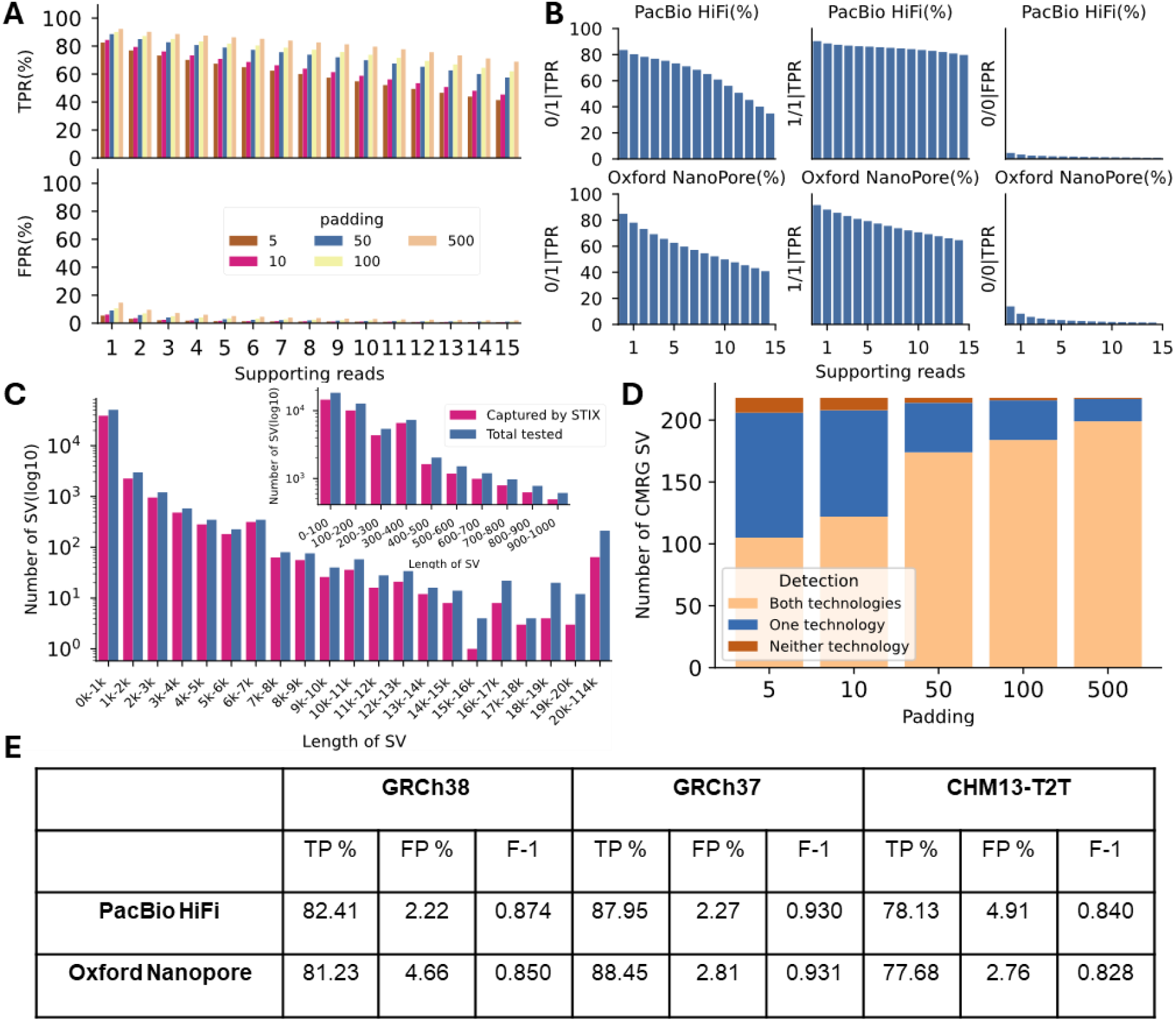
Performance assessment for Germline SV of STIX based on GIAB: Performance metrics for correctly genotyping SVs in different platforms(Pb-HiFi and ONT). All details can be found at **Supplementary Table S2 and S3**. **A)** comparison of Tier1GIAB SV (TPR and FPR) with different support reads(x-axis), padding(color of the bar). **B)** Impact of genotypes on STIX across different platforms. Each sub-figure represents either TPR(first two columns) or FPR(third column) for Pb-HiFi(top row) and ONT(bottom row). X-axis shows the supporting reads. Y-axis is TPR or FPR in percent in each sub-figure. **C)** Assessment of capture rate (y-axis) of STIX for Tier1GIAB SVs across different sizes(x-axis). SVs were grouped by their length. Sub-figure in the right top corner shows the SVs no longer than 1kb with 100bp as bin width. For each group, two bars represent tested(blue) and captured by STIX(red) respectively. **D)** Performance evaluation of STIX for detecting CMRG SV(y-axis) with the increase of padding(x-axis). Colors in each bar represent the detection status(orange:detected in both Pb-HiFi and ONT; blue: detected in either Pb-HiFi or ONT; yellow: not detected in all platforms). **E**) STIX performance with different reference genome and sequencing platforms

For insertions and deletions, the only two SV types reported in the GIAB HG002 benchmark, STIX had a higher TP and FP rate (TPR and FPR) as well as F-1 score in deletions when compared with insertions (**Supplementary Table S1 and Supplementary Fig. S6**). In addition to performing well in genome-wide benchmarks, STIX recovered SVs in regions that are traditionally difficult to resolve. Among the 218 SVs in the Challenging Medical Relevant Genes(CMRG) GIAB benchmark^40^, STIX correctly annotated 89.45%(195) of the SVs based on ONT and 94.04%(205) of SVs based on Pb-HiFi. When the two sequencing technologies were combined, STIX re-identified 216 out of the 218 SVs (99.08%)(**Fig. 2D**). Across the three references, STIX (TPR: 79.7%, FPR:3.27%) also performed similarly to two state-of-art single sample only SV genotypers, Sniffles2^41^ (TPR: 87.2%, FPR: 4.65%) and cuteSV^42^(TPR: 82.3%, FPR: 3.41%) (**Supplementary Table S1**). We further benchmarked STIX with its short-read index that showed clear improvements across the long-read version (**Supplementary Section 5 and Supplementary Figure S9-S15**).

When categorized by zygosity, Pb-HiFi and ONT showed similar TP rates, with heterozygous SVs being lower than homozygous variants (**Fig. 2B**, left). We anticipated this difference due to fewer reads supporting the variant allele in heterozygous locations. The FP rate for Pb-HiFi was about half that of ONT (**Fig. 2B**, right). Among 23,114 heterozygous variants, 5,473 homozygous variants from HG002, and 6,413 wild-type variants from the HG00733 call set(see **Supplementary Section 1**), the TP and FP rates for Pb-HiFi were 87.32% and 2.18%, respectively, while for ONT, they were 87.14% and 4.41%, respectively (**Supplementary Table S4**).

To assess the impact of SV length on the performance, we grouped the GIAB HG002 SVs in different size bins and calculated the recall per 1 kbp bin. As illustrated in **Fig. 2C**, STIX captures 78.33%(average of each length category) of SVs that are less than 10kb (73.09% of SVs less than 15kb). We further stratified those SVs between 50bp-1kbp into 100bp bins as the SV in this size amount to the most number of SV. We observed a good performance in SVs that overlap with Alu (300-400 bp) or LINE elements (6,000-7,000 bp). STIX has limited performance for SVs larger than 15kb but still capture 35.26% of them.

Overall, STIX performed well across both sequencing platforms, regardless of SV zygosity or length. Additionally, STIX can now accurately annotate insertions, a feature not available with the same level of precision as other population annotation methods. We also conducted a benchmark assessment for the short-reads version of STIX to demonstrate the consistency of the new version(**Supplementary Section 5**).

### STIX improves the SV annotation at population scale

While long-read sequencing has well-known advantages for detecting SVs compared to short-read sequencing^1^, as of now, there are no long-read based resources available for annotating SVs. To address this gap, we integrated data from the The 1000 Genomes Project (1KG)^43^ and The Human Pangenome Reference Consortium (HPRC) project^44^ (**Methods and Supplementary Section 3**) to produce a STIX index with 1,108 samples(1,104 for the CHM13-T2T version) from 26 populations and 5 super-populations (**Supplementary Table S5 and Supplementary Fig. S16**) sequenced by Pb-HiFi or ONT at 17x coverage or higher^45,46^.

To assess how accurately STIX can determine the frequency of SVs in a population, we firstly evaluated the consistency of population evidence of 1KG published SV between the long-read index and short-read index with sufficient evidence depth (at least 5) and observed a significant positive correlation(R=0.8, p-value <0.01), as shown in **Fig. 3A**. Moreover, we compared the number of non-reference samples for the same 1KG SVs annotated by the STIX long-read index to those originally reported by the 1KG. There was a significant correlation between the number of samples found by 1KG and STIX that harbored an SV. (R=0.8, p-value<0.001, **Fig. 3B**). About 10% of SVs were more frequent according to 1KG (less than −1 in **Fig. 3C**). These SVs were evenly distributed among SV types (**Supplementary Table S10**) and tended to be smaller (median length 271 bp vs 626 bp for all 1KG SVs). About 20% were more frequent according to STIX (greater than 1 in **Fig. 3C**) and were enriched for inversions (**Supplementary Table S10**), and tended to be longer (median length 6,113 bp). For the remaining SVs, the difference between the STIX and 1KG frequencies was on average 2.2%. Using a Hardy-Weinberg based Gaussian mixture model (**Supplemental Section 6**), we estimated the allele frequency of common (> 10% frequency) SVs that were highly correlated with the 1KG allele frequencies (R=0.73, p-value<0.001, **Fig. 3D**), with a mean difference of 0.6% between our estimated allele frequencies and those reported by 1KG. The STIX long-read results were consistent with STIX short-read results (**Fig. 3E** and **3F**), with the notable exceptions that the short-read database had nearly 2X the number of samples (2,504), the long-read index supports insertions, and the long-read mixture model allele frequency estimations were better calibrated (**Supplementary Fig. S17**).

**Fig. 3.**
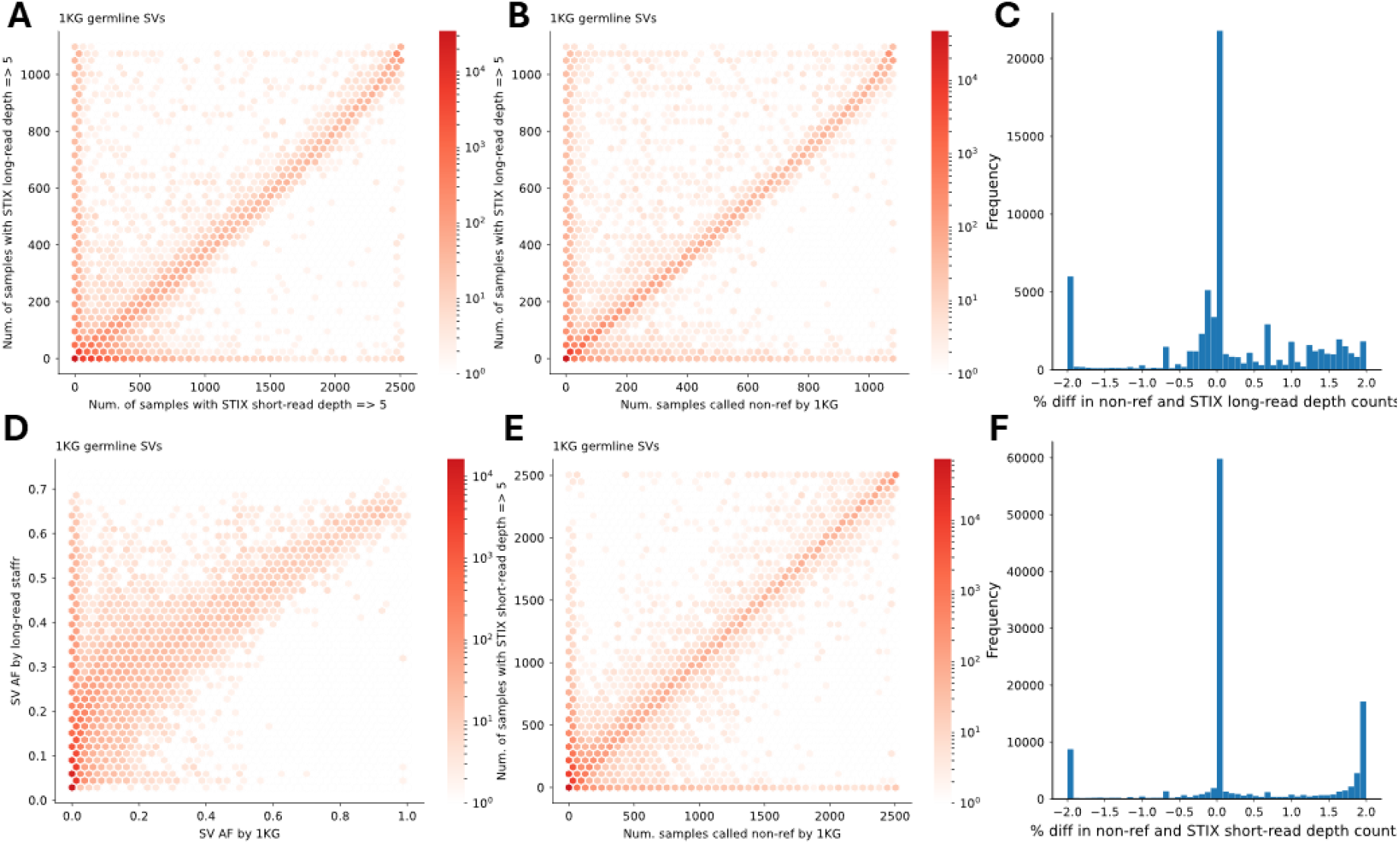
Comprehensive resource of large cohorts: **A)** Comparison between the long-read index and short-read index. **B)** The number of samples with at least 5 STIX long-read hits versus the number of samples assigned a non-reference genotype by 1KG. ***C)*** The distribution of average differences between the STIX long-read and 1KG frequency estimates. **D)** The STIX long-read allele frequency estimated by mixture model versus the allele frequencies published by 1KG. R=0.82. **E)** The number of samples with at least 5 STIX short-read hits versus the number of samples assigned a non-reference genotype by 1KG. R=0.64 **F**) The distribution of average differences between the STIX short-read and 1KG frequency estimates.

STIX offers more comprehensive SV frequency annotation than standard catalogs. To measure this improvement we compared the variant frequency annotations from the 10,847 samples in the gnomAD^24,47^ catalog to the STIX long- and short-read 1KG indices, which had 1,108 and 2,504 samples respectively (the long-read index included 11 PacBio/ONT technical replicates), using the GIAB benchmark sample HG002. **Supplementary Fig. S7** shows the frequency of GIAB SV across the STIX index, which follows the expected distribution. While 99.8% of single nucleotide variants (SNVs) in HG002 appeared in the gnomAD catalog^40^, only 33.4% of structural variants (SVs) were found. In contrast the STIX long-read index annotated 95.9% of the HG002 SVs. Among the gnomAD annotations, there was a notable depletion in the number of insertions. While there was evidence in the STIX long-read index for 94.6% of the HG002 insertions (the short-read index does not support insertions), only 17.3% were in the gnomAD catalog. For deletions, the STIX long- and short-read index annotated 97.8% and 96.8%, respectively, while only 58.6% were in the gnomAD catalog. When considering just the 218 HG002 SVs that overlapped the challaning medically relevant genes (CMRG), the gnomAD catalog only found 69 (31.6%), while the STIX long-read index found all 218, and the STIX-short read index found 97 (41%). These differences are notable considering how many more samples the gnomAD catalog included, and how many of the SVs missing from gnomAD are at high frequency in the 1KG cohort (**Figure 4**).

**Fig. 4:**
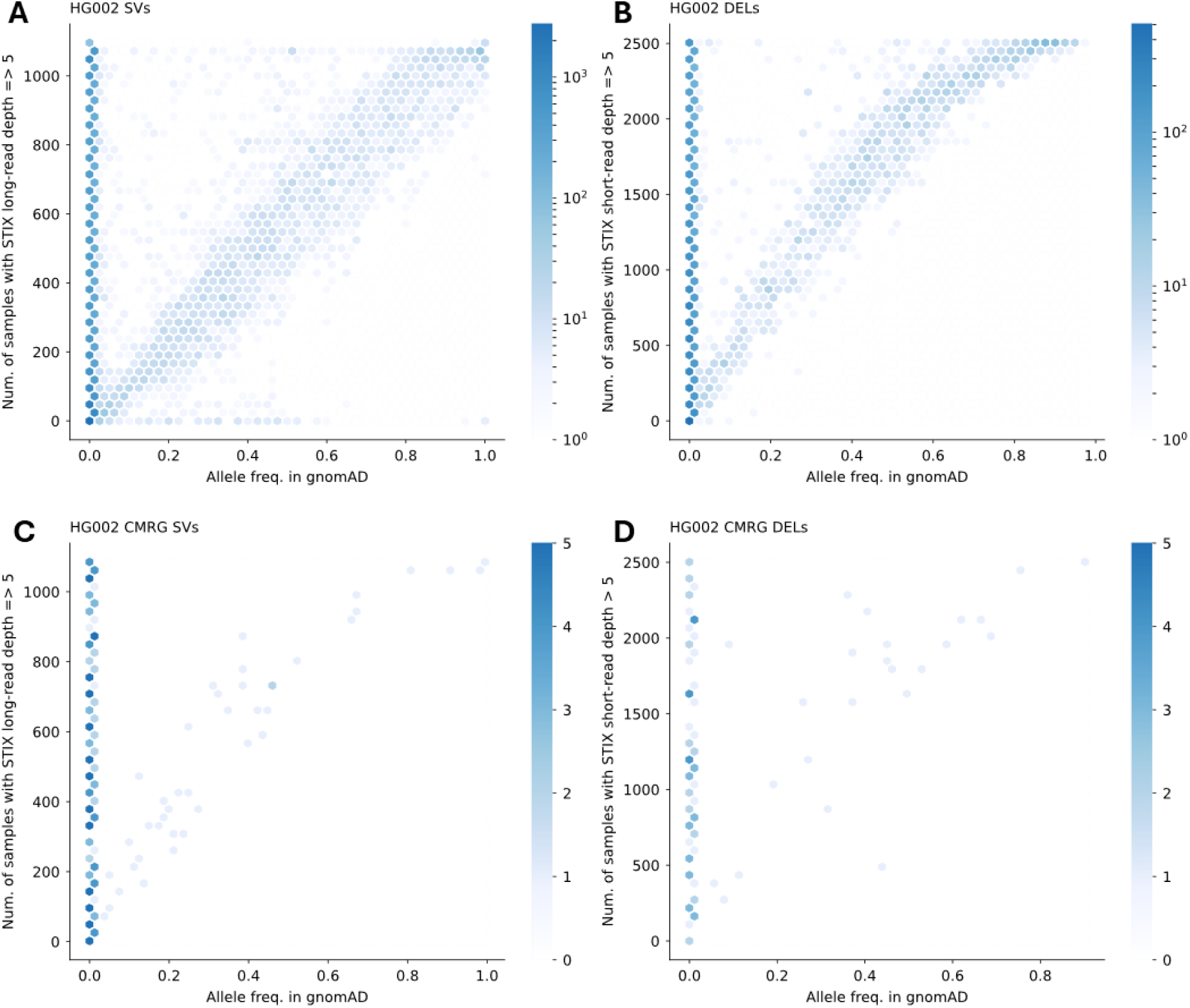
A comparison between frequency estimates annotation of the GIAB HG002 benchmark SVs using the gnomAD catalog and STIX long- and short-read indices. A) The number of samples with at least 5 STIX long-read hits versus the gnomAD allele frequency. B) The number of samples with at least 5 STIX short-read hits versus the gnomAD allele frequency. C and D) The same comparison with the subset of SVs that were in the challaning medically relevant genes (CMRG).

### Improving somatic cancer SV prioritization

SV prioritization, especially in tumor samples, can be complicated by false positives driving the wrong identification of somatic SV^48^. This can be mitigated by improved SV comparison methods, but also by population frequency annotation. For the latter, somatic SV that are not driving the cancer can occur naturally throughout the population and thus might be commonly observed^8,49^. To assess the ability of STIX to improve somatic cancer driver SV detection, we first tested this approach at COLO829/COLO829BL, which are well characterized tumor-normal cell lines^50^. We recently postulated somatic SV for COLO829/COLO829BL^50^ and were interested in how well STIX could identify potentially cancer only SV. We initially started with all SVs (including 29,674 germline SV and 45 somatic SV) from the tumor sample(COLO829). STIX was able to annotate 89.67%(26,649) of the germline SVs. We use 1% AF as the threshold for common variants and found 81.77%(24,302) of the germline COLO829 SVs can be indeed annotated as common in the population and thus were likely non cancer drivers (**Fig. 5A**, purple bars). This is a great result when for example one does not have a normal matched sample at hand at the right quality or quantity to perform e.g. long-read sequencing. We next focused on the previously postulated 45 SV that were identified as somatic for the COLO829 (cancer sample) to further showcase the benefit of STIX^50^. **Fig. 5A**(blue bars) highlights these 45 somatic SV and their proportion of population scale evidence based on STIX. Interestingly, STIX assigned evidence to two of the 45 somatic SV but none of the somatic SV has a higher AF than 1% in our SV index. Thus, highlighting that despite tumor-normal comparison a population annotation such as STIX might further narrow down potential cancer driver mutations compared to likely benign SV.

**Fig. 5:**
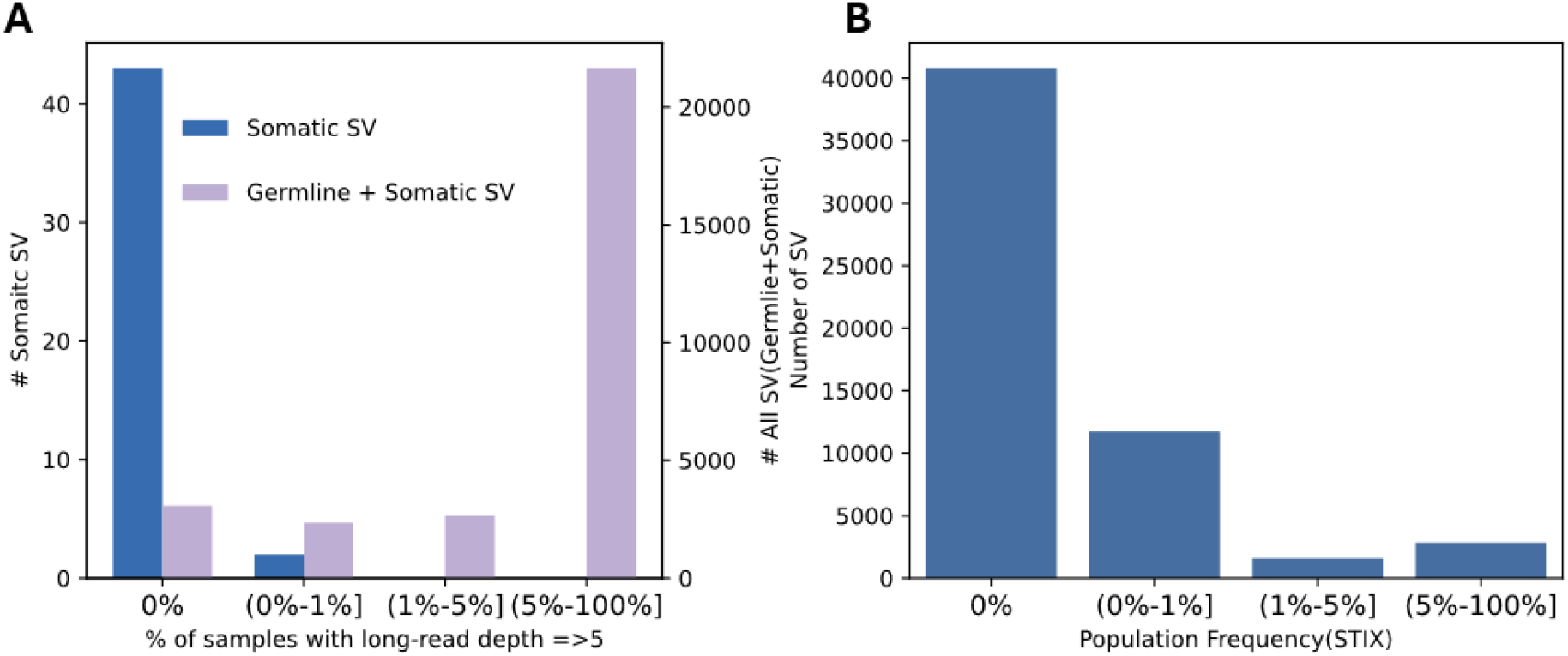
STIX application to cancer structural variation prioritization. **A)** Proportion of evidence distribution of all SV(purple bar,right y-axis) and somatic only SV(blue bar, left y-axis) from COLO829. STIX effectively narrows down the scope of potential pathogenic structural variants (SVs) to approximately 10%, and includes 96% of true somatic SVs, as identified in the matched normal sample COLO829BL. **B)** Proportion of evidence distribution of 46,755 COSMIC SV annotated by STIX. STIX found evidence for 29.01% of these SVs.

Motivated by this observation, we next investigated if we could further annotate and identify SV that are postulated to be somatic cancer mutations, but might actually be common in the population. We use somatic SVs from the Catalogue of Somatic Mutations in Cancer project(COSMIC)^51^ as it summarizes one of the largest cancer datasets. **Fig. 5B** shows the result with respect to the annotatable SV. We annotated 46,755 somatic cancer only SVs using STIX and were able to retrieve population frequencies for 13,564 (29.01%). Among them, we identified 3,563 SV (26.27%) as common in the population (>1% AF) indicating their potential non-pathogenic role. These SVs could also represent non-cancer driving somatic SVs that were accumulated during cancer initiation and progression and thus be potentially harmless^52,53^. However, some SV could potentially also represent several falsely identified somatic SV in COSMIC, which is hard to determine. The remaining 10,001 (73.73%) SV showed a low proportion of evidence(<1%) as may be expected from somatic benign or cancer SV postulated by COSMIC.

Across both examples, STIX showed to be important in further prioritization of cancer vs. benign somatic SV. This is obviously just one of the potential important use cases of STIX for the prioritization of SV across many human genetic diseases.

### Exploring mosaic SVs across population level

Following the identification of potentially common somatic SVs in the population, explored STIX’s ability to identify mosaic SVs (i.e., those with a low variant allele fraction (VAF)). To benchmark STIX’s ability to assess and annotate mosaic SVs, we created STIX indexes from read sets sampled at different rates ranging from 0.05 to 0.45 from HG002 and HG00733. For each index, we searched for evidence of the HG002 benchmark SVs (**see methods, Supplementary Section 2**). Overall, STIX had an average of 90.61% TP for deletions and 64.59% TP for insertions(**Fig. 6A**). We speculate that the random choice of reads favor the more frequent shorter read length in the distribution of HG002 and thus reduce our performance on insertions. Similar to the germline ONT benchmark, we observed an impact of the padding parameter on the identification ability of STIX and did not observe a bias towards SV length with 95.27% of all SVs captured across the different lengths (**Fig. 6B**). Among the CMRG regions (**Supplementary Fig. S18**), STIX captured 182 SVs (83.49%) despite the SV being present at lower VAF. Overall, STIX showed great performance in annotating mosaic SV despite a greater challenge of lower number of reads present per SV.

**Fig. 6.**
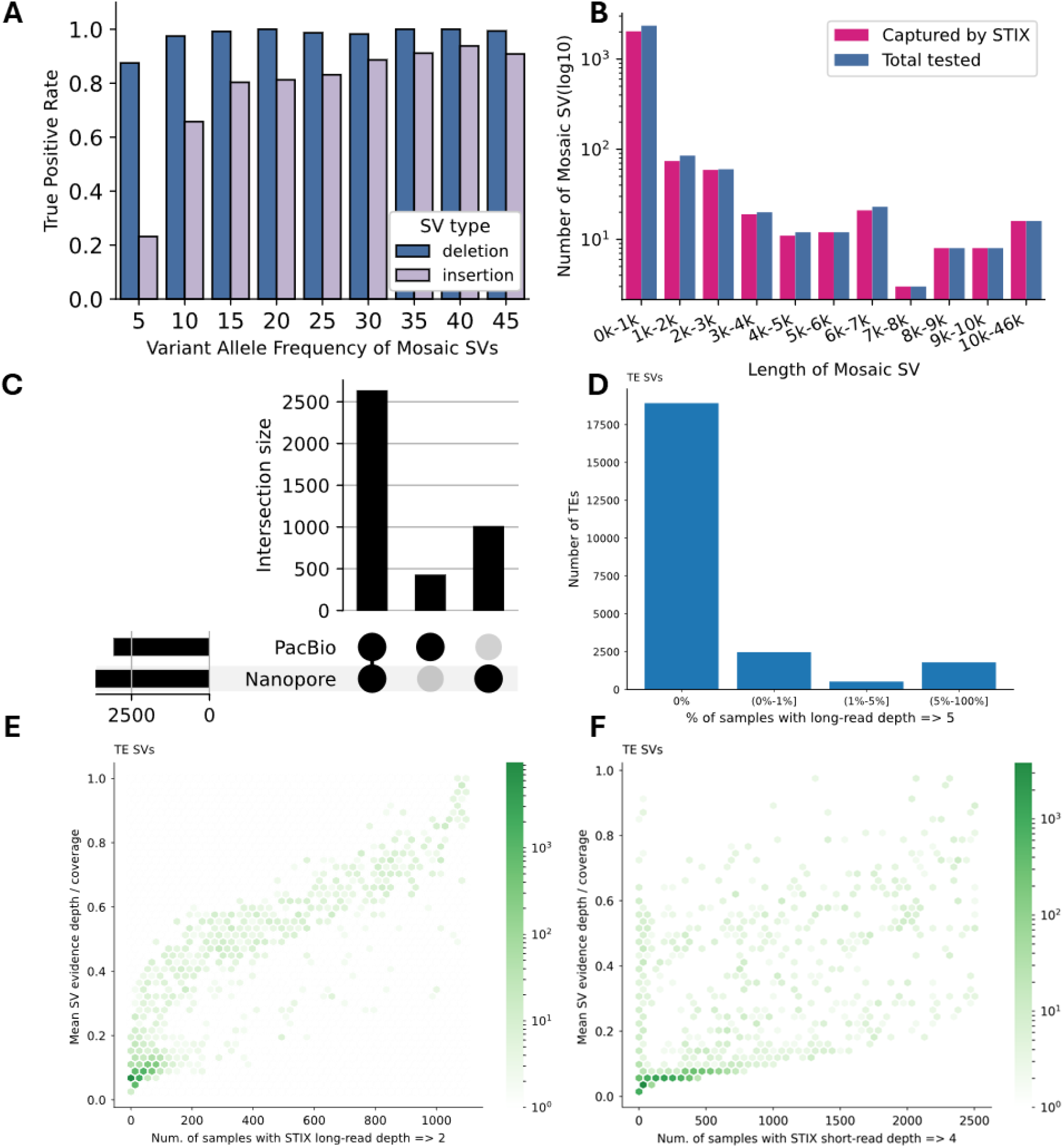
Investigation of mosaic SV at population level: **A)** True positive rate of mosaic SVs. SVs were spiked in with different fractions(x-axis) to simulate various VAF. Each VAF has two bars representing deletions(blue) and insertions(yellow) respectively. **B)** The impact of SV length. Mosaic SVs were classified using 1kb bin size(x-axis). y axis represents the number of SVs with log 10 scale. The blue bars represent SVs total assessed, red bars represent SVs captured by STIX. **C)** the comparison of 6 samples sequenced by both Pb-HiFi and ONT. SVs were grouped by their type(DEL,DUP, and INV) and the existence of two platforms. **D)** Proportion of evidence distribution of mosaic TEs that was annotated by STIX. **E-F)** Distribution of average variant allele frequency of a SV in samples(y-axis) versus number of samples that harbor the SV(x-axis). Variant allele frequency of a SV in a sample was calculated by the supporting reads / sample mean coverage. **E:** long-read index, **F:** short-read index

Next investigated if a mosaic SV that was induced by a germline insertion identified in our recent work^41^, was more common in the population than expected. In our previous work, we identified and validated a 6% VAF deletion that was due to an apparently common germline insertion of an ALU-Y element near an existing ALU-Y^41^. These two ALU-Y seemingly recombined in a certain number of cells and thus lead to a deletion signal of ∼6% VAF. Over PCR and sanger validation we could confirm these variants, but also saw emerging bands in other brain samples that we tested. Thus, we speculate that since the germline insertion is common in the population that the recombinant mosaic allele might also be common in the population. We used STIX to annotate both events reported and validated previously. STIX reported the ALU-Y insertion in around 48.10% (533 samples) of the samples clearly confirming our suspicion that this is a common germline insertion in the population. The resulting recombinant between the two ALU elements was found at 0.2% proportion of evidence in the population(2 samples). It’s noteworthy that if we reduce the read threshold to 3, given the overall coverage of 15-20x across the samples, we identified 23 samples(2.08% proportion of evidence). This indicates that while this is a somatic/mosaic variant it is also commonly independently gained through the population.

This observation spiked our interest in what other repeat recombinants might be commonly shared in the population despite being independently gained throughout the population. To further extend the search for common, but mosaic alleles in individuals we next studied a recently reported set of mosaic transposable elements^54^. First, we focused on six samples where we had matched Pb-HiFi and ONT data available in the STIX index. Thus, if we would observe the consistency of these alleles across both technologies, it would strongly indicate that some of these postulated mosaic SVs are shared in the population. We collected 18,331 mosaic SVs (3,551 deletion, 1,224 duplication and 13,348 inversion) from the CELL paper and removed the duplicates as they were originally discovered at per-read level. We identified 9.28% SVs being annotatable in at least one sample. Among them, 64.9% SV were identified both in Pb-HiFi and ONT indicating the high-consistency across sequencing technologies(**Fig. 6C**). Interestingly, there are 84.7% deletions and 69.0% duplications having support by both Pb-HiFi and ONT, while only 6.32% inversions were observed in both technologies. While there are many more inversions reported by the previous study, we were curious on why there is such a discrepancy. Given this observation, we investigated ONT and Pb-HiFi based indexes separately. Over manual inspection (**Supplementary Fig. S8**) we speculate that these inversions are false positives in the previous study caused by chimeras found in ONT data. We had reported something similar for inversions before^41^ but didn’t expect to find this artifact in a published SV call set. Interestingly, these inversions are typically smaller with a median size of 573 bp, which further matches our expectation of likely artificial chimeras at low frequency. Thus, we concluded that many of these inversions are false positive SV calls in the previous CELL publication^54^, which further explains their high number of events (2.7 fold more inversions than deletions and insertions) Next, we focus on mosaic TEs that are presented as deletions as they show high consistency. Overall, 23,703 SVs(deletions) were included. Among them we can identify 6,100 (25.7%) SVs in the healthy population and 2,324 (10.1%) are even common(>1% population evidence). This suggests that mosaic SVs are more often independently gained than we might expect. For those SVs that are shown positive signals at population level, the majority of them exist at low VAF status while the others tend to present in the population status (**Fig. 6D**). Despite those SVs with low VAF and low proportion of evidence in the population(**left bottom part in the Fig. 6E**), we observed a linear pattern showing a clear correlation between VAF and the proportion of evidence in the population (**Fig. 6E**). This indicates that some of the postulated mosaic repeat recombinants rose to germline SV. This observation is also supported by the short-reads version of STIX (**Fig. 6F**).

## Discussion

In this work, we present a long-reads based annotation approach by extending STIX to long-reads datasets. This new version now also supports insertions and can utilize either Pb-HiFi or ONT data. We validated the performance of STIX for long-reads with the GIAB benchmark(HG002 sample) and assess its specificity using a negative control sample(HG00733). We found that the overall true positive rate (TPR) is 81.82% and false positive rate (FPR) is 3.44%. We observed higher performance of STIX and other two state of the art SV genotypers on the GRCh37 version benchmark as it is better established than the GRCh38 and CHM13-T2T benchmark. According to our experiment, STIX shows robust performance with SVs with different length and genotype. We further tested SITX in SVs that are located in challenging medical relevant genes(CMRG SVs). We achieved 89.45% TPR in both sequencing platforms (Pb-HiFi and ONT) and 99.08% TPR when considering at least one sequencing platform. Furthermore, it is also important to highlight its future readiness as the index is ready to be extended, can include meta information (e.g. ethnicities, diseases background etc) and does not rely on a SV calling pipeline that ever needs to be updated. Thus, STIX demonstrates a clear advantage when annotating SV with population frequencies.

Variant annotation and thus prioritization are of utmost importance to identify potential pathogenic variants for the medical and biological sciences^55,56^. The principle still holds that common variants in the population for the most part are not pathogenic^19^. Therefore the identification of rare variants based on the population represents still one of the best practices to rank or prioritize variants. Over the past decade multiple advancements in SNV annotation have been made, which greatly improved medical and clinical studies^57–60^. These advances have been mainly achieved by short-reads as long-reads so far have been cost prohibitive to build up significant annotation resources. Thus, despite the fact that long-reads improve SV detection, short reads have been utilized to annotate SV^24^. This has led to the issue that many SVs are not annotatable and thus the prioritization was not possible. Therefore the identification of pathogenic SV has been hindered. This is demonstrable by gnomAD only being able to annotate 33.5% of the HG002 GIAB SV catalog despite the sample likely being present in gnomAD as the SNV are annotatable by more than 99.7%. To overcome this, we have extended STIX to incorporate long-read information for SV annotation.

With this we are now able to annotate and thus prioritize SV. Despite the significantly smaller indexed population size of 1,108 genomes, STIX was able to demonstrate significant concordance with large short-read based catalogs for SV that were annotatable in both data sets. This together with low false positives rate really improves the SV annotation. Another important aspect about STIX is also the accuracy it annotates variants avoiding the commonly used reciprocal overlap of 50% or 70%^24^. It is important to also discuss the behavior of STIX when annotating false positive SV themselves. As highlighted with inversions, this typically results in high population evidence annotations from STIX. This typically would discard these false positive SV or artifacts from subsequent studies. Of note this is in contrast to the FPR of STIX, which represents the annotation of a similar but different, close by SV. In this work we have carefully benchmarked and concluded that actually a 100bp padding and min 5 reads are sufficient to accurately annotate SV and avoid FPR with close by SV.

We were able to demonstrate STIX ability and utility in cancer data sets and mosaic SV call sets. For cancer we identified 3,563 SV that are reported in COSMIC^51^, but are actually common (>1% proportion of population evidence) in the population. While it remains unclear if these are errors in COSMIC or maybe common genome instabilities, it shows that STIX is an important method to prioritize cancer mutations or avoid potential common SV. Thus, also speeding up the prioritization of SV for cancer studies. Based on our experiment, STIX can narrow down the scope of potential pathogenic SV to approximately 10% and retain 94% of true somatic SVs. This demonstrates clearly that STIX is useful in the case of tumor only or even paired sample analysis. To investigate the potential for common mosaic SV, we have annotated a previous validated mosaic deletion, which was caused by an Alu-Y repeat recombinant ^41^. STIX was able to annotate both events and indeed showed that the germline Alu-Y insertion at this location was highly common in the population (48.10% of the individuals). In addition, the resulting mosaic deletion was still detected in two samples (increases to 23 when reducing the thresholds) despite the general lower coverage of the indexed data sets (15-25x). We expanded this study by investigating somatic repeat recombinations previously published. Here STIX was able to find 10.1% of the SVs are commonly shared(>1%) within different individuals of our index. Thus highlighting that mosaic SV despite being rare in an individual can be independently gained thought the population by likely common genome instabilities (ie. repeat recombination sites). This result is significant as it demonstrates the plausibility of the 3,563 COSMIC somatic SV that are commonly shared not being false positive, but rather benign variants. This is consistent with previous studies^52,53,61,62^. Thus, clearly highlighting the importance of STIX to be used to annotate SV to streamline pathogenic SV detection itself. This finding further highlights the need to study and understand mosaic SV further as some fraction seems to be commonly shared in the population and even arise to higher zygosity.

Lastly, we compared the new long-read version of STIX presented in this paper with the short-read version, finding a high level of consistency between them. This demonstrates STIX’s robustness across both long-read and short-read data, paving the way for joint SV annotation using long-read and short-read datasets simultaneously. A gaussian mixture model was developed to provide the estimation of the population allele frequency of query SVs based on the amount of evidence (the number of alignments) that support the presence of a query SV in a sample(and population of samples) that was returned by STIX. We tested this model with both long-read and short-read indices and observed a high correlation with the original allele frequency from 1KG. As expected, the correlation was higher with long-reads (**Supplementary Fig. S17**) given its lower noise profile. Since the model was based on the Hardy-Weinberg equilibrium, it struggles to classify SVs with allele frequencies greater than 0.7, and most samples have two copies of the variant.

Overall, STIX has shown to be a versatile and accurate SV annotation methodology that represents a significant improvement over other resources. Its indexing of raw reads makes it updatable and future ready for improvements in SV identification over the next few years.

## Methods

### Extending STIX to long-reads datasets

STIX extends the ability to extract, index, and search all types of SV signals from datasets sequenced by ONT and Pb-HiFi platforms (**Figure 1A**). To accomplish this, we developed a new program named excord-lr, which is the long-read version of excord^26^ to extract signals from long-reads datasets. This program was written in the Rust programming language. Precompiled binaries are available under releases in its GitHub repository. Briefly, the Excord-lr program takes the BAM file as input and extracts SV signals from both the CIGAR value and SA tags. It incorporates a variety of empirically-based filters to minimize false positive signals. This includes 1) controlling the maximum number of supplementary alignments for a single read, and 2) limiting the proportion of overlap between two supplementary alignments. A function was also embedded in Excord-lr in order to extract insertions with different situations. The output is compatible with short-version, which generates a file with BEDPE format. Excord-lr also supports running with multiple threads to accelerate the speed. Debug information and reads id of a SV signal are also available under the debug mode. Excord-lr is specifically designed to manage the distinct features of long-read sequencing datasets and to ensure they are compatible with further searching and indexing processes.

One of the major updates in STIX is supporting the extracting, indexing and searching of the insertions with different lengths. To accomplish this, STIX uses excord to extract three types of insertions encompassed in long-read sequencing datasets according to its length,mapping status, and the fields present in the BAM file.

1. Shorter insertions will be encoded as “I” tags in CIGAR values.
2. Large insertion will be encoded as supplementary alignment in the SA tag. But the insertion is shorter than the read length, but can not be presented in CIGAR value. It may generate a primary alignment and a supplementary alignment. Each alignment will have a soft-clip or hard-clip on the left and right respectively.
3. When the large insertion is longer than read length, it will only result in a primary alignment with one large soft-clip or hard-clip only.

Taking into account those conditions, excord-lr implies three different approaches to extract insertion signals. For the insertions in CIGAR value, it will extract the position of insertion point as well as the length of insertion. For insertions present in SA tag that generate two alignment, excord-lr will extract them by using the criteria as follows:

1. must have one primary alignment and one supplementary alignment.
2. Each alignment must have a soft-clip or hard-clip(no less than 1kb by default).
3. two alignments should overlap to each other.
4. both alignment should have the same chromosome name.
5. For insertions that only generate one primary alignment, excord-lr extract them by including the following filters:

a. only one primary alignment, no supplementary alignment.
b. primary alignment must have soft-clip or hard clip more than 1kb.

Excord-lr encodes insertions as 0 base pair intervals in BEDPE format. The insert point is presented at the end of the left region and the start of the right region. In other words, the end position of the left region will equal the start position of the right region if the record encodes an insertion. This 0 base pair interval will be used to STIX searching step to distinguish insertions from other types of SVs. if the insertions are accompanied with length information(insertions that are extracted from CIGAR value), the length will be encoded as the length of the right region, otherwise, the start position is equal to the end position in the right region.

STIX kept the same steps to index SV for signals from long-reads datasets with the short-reads datasets, as described^26^. Briefly, Giggle, a fast genomics search engine, is used as a dependency to index SV signals^36^. The output bed file from excord-lr will be sorted according to the position of the left and right region and be compressed with gzip. For example:

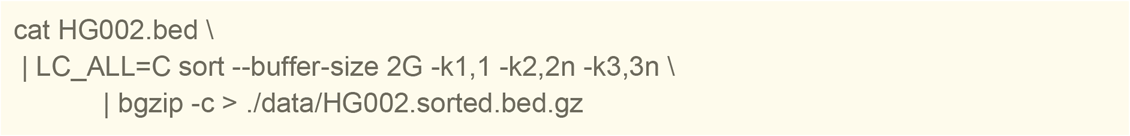

Giggle index then can be built using command as following:

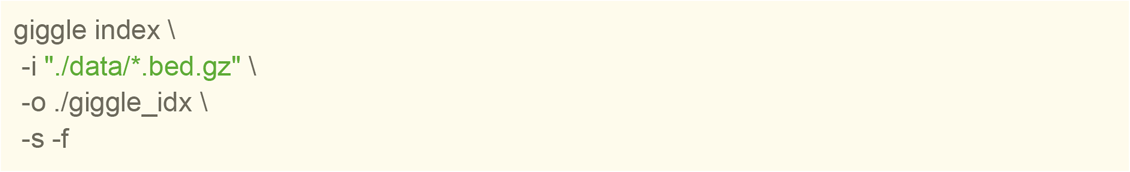

STIX index will be created based on top of sorted bed file, giggle index, and metadata.

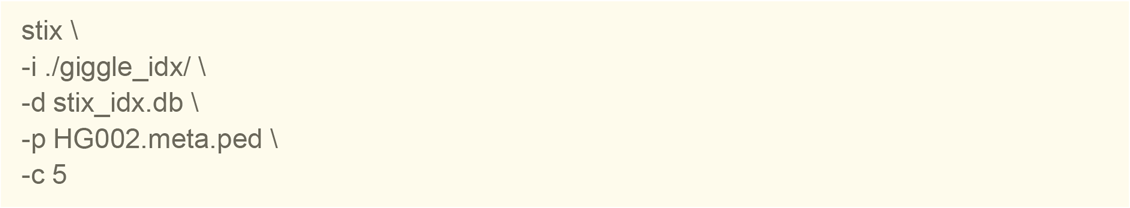

STIX requires a file that describes sample-related information, which can be specified with the-p option. The -c option is used to indicate the column of the file names.

The search step in the current STIX version is mainly inherited from the old version but with a few new options to enhance its ability. Concisely, STIX can take a single region pair or a VCF file as input. It can also accept a tabular format as input in the latest version in case users find it difficult to generate a VCF when they want to query more than one region pair.

When searching for an insertion. Users need to provide two base pair intervals for left region and right region respectively as well as an additional parameter (-L) to specify the length of the insertion. The steps for how STIX searches for an insertion are briefly outlined as follows:

1. searching the potential records that overlap with the query regions.
2. For each record, mark them as an insertion record if the end position of the left region equals the start position at the right region and the chromosomes of both regions are identical.
3. Compare the insertion point between the query and each result, report a hit if they are identical, otherwise report a non-hit.
4. If the length of query insertion is provided by the user, STIX will try to further compare the length between the query insertion and the potential insertions from the index. A relative error will be calculated to describe the similarity of the length between the query and we set 0.2 as default cutoff.
5. If the user does not provide the length information for query insertion. STIX will search the potential insertions located in the padding region.

There are examples for running STIX in different search mode:

1. Single query:

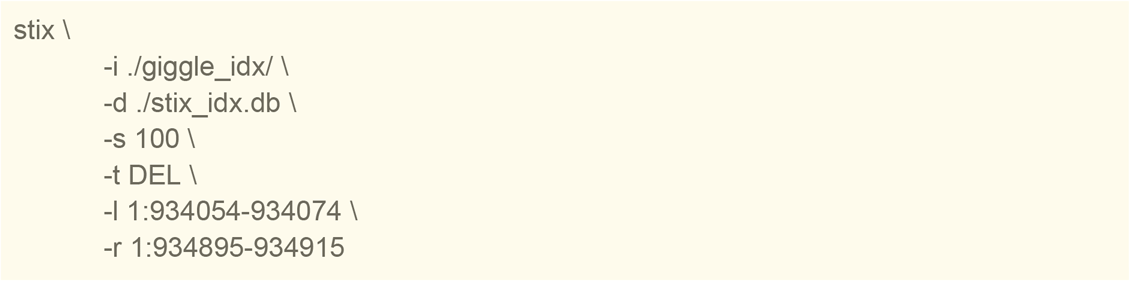
2. Batch query using VCF as input:

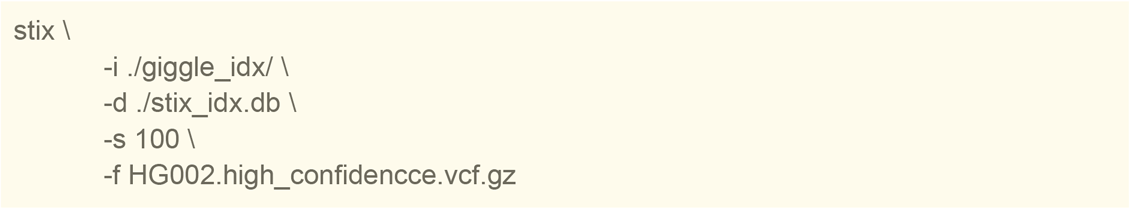
3. Batch query using table as input:

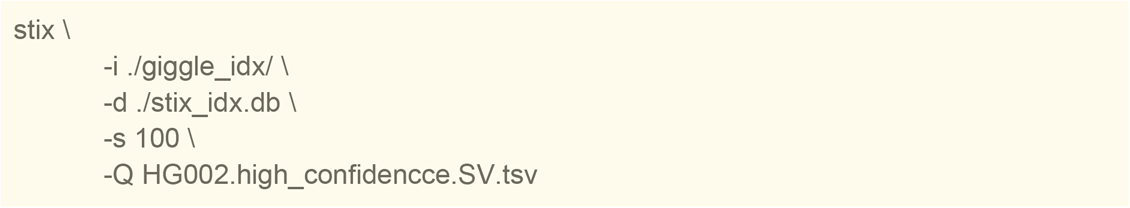

Additionally, we introduced a sharding mode to facilitate indexing of large datasets. This has a wrapper script for generating separate Giggle indexes and an -B option in STIX, which accepts a two-column TSV file. Each row in the file specifies a Giggle index and the corresponding STIX index.

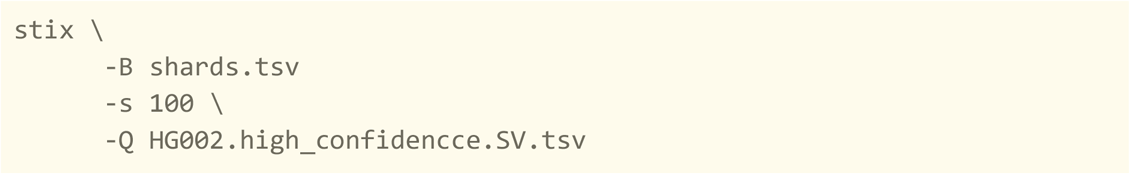

### Data collection and preprocessing

The original links of all public datasets can be found at **Supplementary Table S1.** Briefly,the SVs of HG002 in the Variant Call Format were downloaded from the Genome In A Bottle project(GIAB). The accompanying high-confidence regions were also downloaded from the same path. The GRCh37 version of the SV callset of HG00733 was collected from the Truvari project^25^. We generated the GRCh38 version and CHM13-T2T version callset for HG00733 using an assembly-based pipeline which replicated the GIAB TR process for variant mapping^63^. We initially collected two bam files of HG002 which were sequenced by Pb-HiFi and ONT respectively, and a bam file of HG00733 that sequenced by ONT. All bam files were aligned to GRCh37 reference and were remapped to GRCh38 reference and CHM13-T2T reference respectively using samtools and minimap2. The bam files for long-reads index were downloaded from The 1000 Genomes Project and The Human Pangenome Reference Consortium project. Raw reads were extracted from HRPC bam files and realigned to the GRCh38 and the CHM13-T2T reference using minimap2. Callsets from COLO829 and COLO829BL were collected from https://zenodo.org/records/10819636. CMRG SVs callset were downloaded from the GIAB project and intersected with the high-confidence regions. IDs for each variant were added using perl -pe ‘if(!/^#/){$c++; $_=∼s/\t\./\tGRCh38_CMRG_$c/}’ to make it meet the input requirement of SVAFotate(0.0.1). High-quality SVs from high-coverage short-reads dataset in the 1KG project were collected from this paper^64^. The raw vcf was downloaded from 1KG FTP site and retained samples that only overlap with STIX LR index by using bcftools view -S sample_id.unique.txt --force-samples -o 1KGP.subset.vcf 1KGP_3202.gatksv_svtools_novelins.freeze_V3.wAF.vcf.gz. Mosaic transposable elements (TEs) were collected as requested from the authors^54^. The raw format is tabular format and was transformed into a 5-columns tabular format (left-region \t right-region \t length \t svtype\t ID) using python. The projection of svtype is listed below: “del” --> “DEL”, “dupl” --> “DUP”, “Inter” --> “BND”, “inv” --> “INV”. Raw fastq file of TP10Plus was downloaded from under SRA(PRJNA636606) and remapped to GRCh38 using minimap2(2.26-r1175).

### Benchmarking for germline SV

Raw SV signals were extracted using excord-lr(version 0.1.17) and build index by using STIX for HG002 BAM files for both Pb-HiFi and ONT platform respectively as described above and the previous paper^26^ and https://github.com/ryanlayer/stix. To collect high-quality HG002 SVs and ensure the consistent existence of those SVs in HG002 bam files, we only consider the SVs located in the high-confidence regions provided by GIAB. SVs that were larger than 50bp were used for downstream analysis. HG00733 specific SVs were generated by excluding the common SVs from HG002 callset by using SURVIVOR(1.0.7)^65^ with parameters ‘1000 1 1 1 0 50’. This merges the VCF files with 1000bp padding enforcing the same type and strand for variants larger than 50bp. Based on final VCFs for HG002 and HG00733, 5-columns tables were generated for STIX one by one query mode. STIX took SVs from HG002 and HG00733 as input and query each SV on the HG002 index using command:

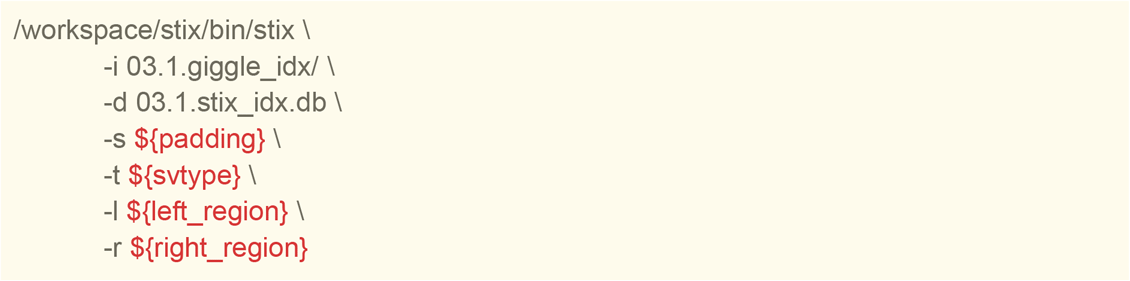

We use different padding: 5, 10, 50, 100, and 500 to test the performance and consider the minimal supporting reads from 1 to 15. We consider the number of hits in HG002 SVs as true positives and non-hits as false negatives, the number of hits in HG00733 as false positives and non-hits as true negatives. Other statistics can be calculated using the following equations:

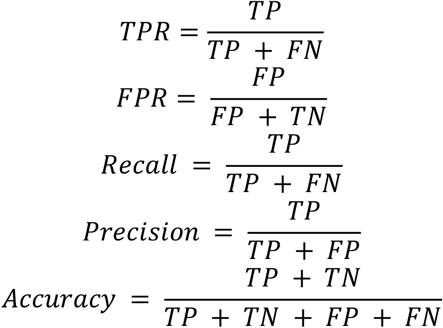

Run Sniffles2^41^ (2.2) genotyping mode:

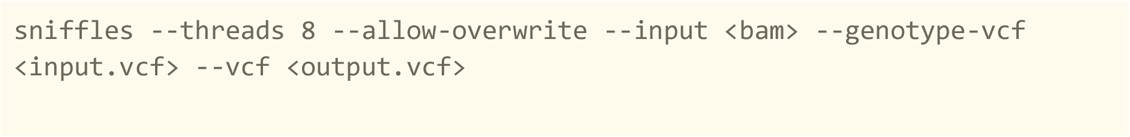

Run cutesv^42^(2.2.1) genotyping mode:

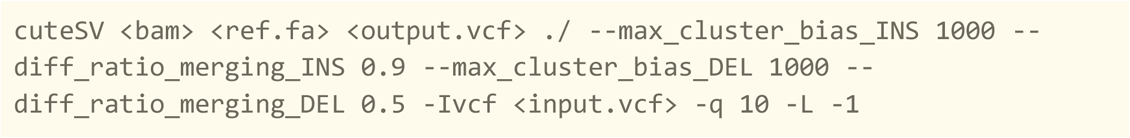

### Benchmarking for mosaic SV

We have developed a framework for generating in-silico mosaic structural variations (SVs) with varying variant allele frequencies, in addition to creating corresponding callsets. Briefly, mixed BAM files that are used in the mosaic benchmark were generated by injecting raw reads from HG002 into HG00733 bam. We used samtools to subset the HG002 bam with the -f parameter to specify the percentage of downsampling, and merged the subset BAM with the HG00733 bam. Finally, the merged bam files were sorted and indexed. Sequencing depths of the BAM files are estimated using mosdepth^66^. SV callsets from HG002, which are used in the germline benchmark, were re-genotyped in the mixed BAM files using Sniffles (version 2.2) in force-call mode with the parameters --input mixture.resort.bam --genotype-vcf HG002.vcf -- vcf mixture_regenotype.vcf --allow-overwrite. Only the SVs that were reported with supporting reads were retained for downstream analysis. Variant allele fraction(VAF) were calculated based on the AD and DP in the output VCF file. Raw SV signals were extracted using excord-lr and the STIX index was built based on top of the sorted bed files with command listed in the Methods.

### Build large index on long-reads datasets

We gathered publicly available long-read sequencing datasets from healthy individuals. In summary, we obtained 100 datasets generated using the ONT platform from the 1000 Genome Project. These datasets were already aligned to the GRCh38/CHM13-T2T reference genome using minimap2 ^67^. Additionally, we acquired 100 datasets (in unaligned BAM format) generated using the Pb-HiFi platform from The Human Pangenome Reference Consortium project. We aligned those dataset to GRCh38 and CHM13-T2T reference with the following command: ‘minimap2 -ax map-hifi <GRCh38.fa> <sample.fq.gz> | samtools view -b > aln.bam’. We downloaded 908 datasets from the 1000 Genome Project (Vienna project) with GRCh38 and CHM13-T2T reference individually. The depth of coverage for each dataset was calculated using mosdepth^66^ with the following parameters:’ -b 1000 -x -t 8 -Q 20 --no-per-base. The ethnicities of those individuals were annotated by manual curation. The raw SV signals were extracted by excord-lr and the sharded giggle indices and STIX indices were created as described above.

### Annotate SVs based on short-reads datasets

We use SVAFotate^24^ to annotate SVs that are used in this study. SVAfotate software (version 0.0.1) was installed according to their instruction^24^. The corresponding libraries were downloaded from the same repository. We downloaded the SVs from Challenge Medical Relevant Genes from GIAB with the filename HG002_GRCh38_difficult_medical_gene_SV_benchmark_v0.01.vcf.gz. We selected the SVs located in high-quality regions by bedtools(v2.30.0) using command:

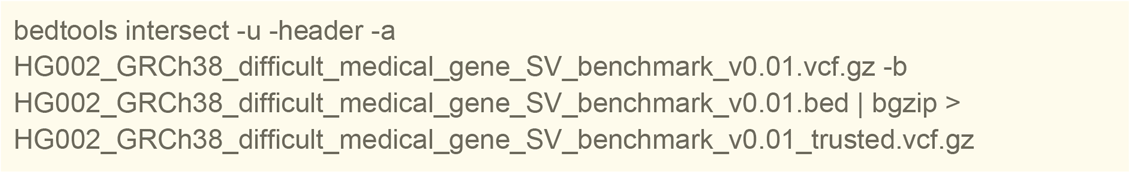

Unique IDs were manually added for each SVs since the SVAFotate needed them to make the right output. The annotate command used in this study is listed below:

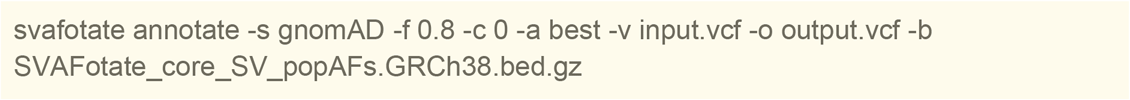

### Analysis of Mosaic Repeat Elements

Raw data of TP10plus were downloaded from the SRA database and converted to fastq format using SRAtools. We next align the raw data against the GRCh38 reference using minimap2 with parameter “ -t 8 -ax map-ont -Y”. Then use samtools to convert the SAM file to BAM format. Excord-lr was used to extract SV signals from aligned bam and STIX index was created by using the aligned BAM file. Mosaic repeat elements locus were downloaded from the supplementary materials of previous published work^54^. We further convert the format for STIX searching. In summary, for each original file, we extract the left and right breakpoints. The duplicated records after rearrangement of left and right breakpoint were removed. We projected the SV types from .out file to STIX accepted format by the following rules: del → DEL; dupl → DUP; Inter → BND; inv → INV. a 5-column table was generated for each .out file and further used for STIX annotation. SVs that were not located in the GIAB high-confidence regions were excluded to avoid the noise.

### Hardy-Weinberg mixture model for estimated an SVs alternate allele frequency

STIX returns the amount of evidence (the number of alignments) that support the presence of a query SV in a sample. While *evidence depth* is useful for reasoning about an SV’s role in a trait, it is not a standard population genetic metric and its unfamiliarity can limit its usefulness. The ideal population genetic metric for estimating a variant’s impact is population allele frequency, or the proportion of haplotypes in the population that harbor the variant. The primary difference between *evidence depth* and *allele frequency* is that the former is a binary reporter–does a sample have evidence or not–while the latter sorts samples with evidence into heterozygous or homozygous states. To quantify these additional states we leverage two observations. First, is that when STIX is performed across a population, the distribution of evidence depths for many SVs has 3 rough modes that correspond to the samples that have zero, one, and two copies of the variant. Second is that past analyses indicate that 86% of SVs are in Hardy-Weinberg equilibrium^22^.

From these observations we developed STAFFR (structural variant allele frequency finder, https://github.com/behzodcu/staffr), a probabilistic generative mixture model that embodies the Hardy-Weinberg equilibrium (*p*^2^ + 2*pq* + *q*^2^ = 1), which can be used to estimate the alternate allele frequency *q* from STIX evidence. Our model interprets variabilities in sequencing data, attributing different levels of evidence depths to specific genotypes: low or very low depths correspond to a homozygous reference genotype where the structural variant is absent, medium depths correspond to heterozygosity with one copy of the variant present, and high depths correspond to a homozygous recessive genotype with two copies of the variant present. This probabilistic generative mixture model is based on modeling variation in evidence using Normal or Gaussian distributions, which have a probability density function of:

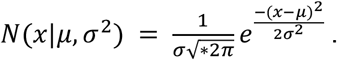

Here μ and σ^2^ denote the sample mean and variance of the variation in evidence across individuals. In general, such mixture models are defined by a collection of several Gaussian distributions, each with a distinct mean and variance, which jointly represent the likelihood of the observed data. The likelihood of each particular observation depends on its total probability across each distribution or mode in the model, which has the form:

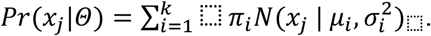

Here Θ = {π, μ, σ} where π_*i*_, μ_*i*_, and σ^2^_i_□ are the probability of generating a value from the *i*th mode, that mode’s mean and its variance, respectively.

Integrating the Hardy-Weinberg equilibrium into the framework of a mixture model allows for the incorporation of genetic principles directly into the model’s estimation. Specifically, Hardy-Weinberg equilibrium defines a mathematical relationship that governs the proportions of genotypes in a population based on allele frequencies and their relative levels of evidence. According to the equilibrium, the frequencies of homozygous reference, heterozygous, and homozygous alternate genotypes occur as *p*^2^, 2*pq*, and *q*^2^ respectively, with *p* and *q* representing the frequencies of the two alleles. By constraining the model’s estimation to conform to these expected genotype frequencies, a modified mixture model embodies the foundational concepts of population genetics, ensuring that the estimated genotype frequencies are consistent. In the above mixture model, we formalize this notion by associating each genotype with the proportion of evidence depths there are, i.e., with their weight π = {*p*^2^, 2*pq*, *q*^2^}.

Furthermore, a structural variant in Hardy-Weinberg equilibrium implies a constraint on the model’s estimated evidence depths based on the presence of one or two copies of the structural variant. As μ represents the mean evidence depth associated with a single copy of the variant under the model, the implied mean for the mode representing two copies of the variant will be twice the mean of one copy, μ_3_ = 2μ_2_. And, considering that the reference population exhibits no evidence of structural variant presence, we replace the first mode with a simple proportion π_1_ = α of data showing no variant, i.e. the proportion of zeros.

Putting these together, the total the likelihood of the observed data being generated from any of the modes with these constraints, is given as:

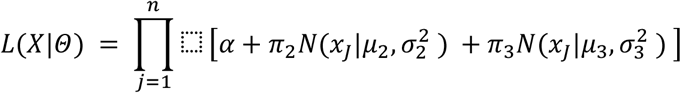

with log-likelihood

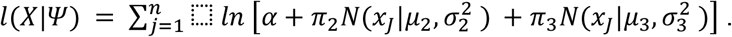

We identify divergences by the observed data from the Hardy-Weinberg equilibrium using a null model approach^68^. In our null (non-HW) model, *q* = 0 and the noise from sampling the reference genome is modeled as Ψ = {α, λ}, where α is the proportion of zero evidence data points (as in the HW model) and non-zero evidence values are modeled by a geometric tail with an expected value λ:

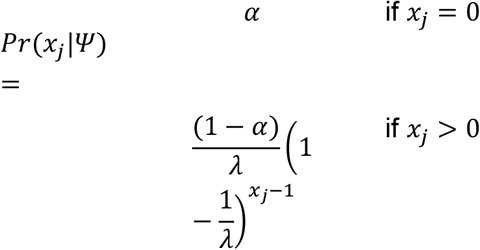

In the null (non-HW) model, the likelihood of the observed data is given as:

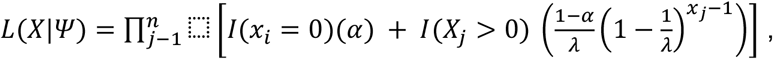

where we use indicator variables *I* to split the values of *x*_*j*_ between zero and non-zero. The log-likelihood is then

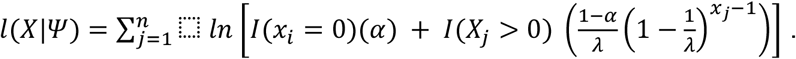

The free parameters Θ and Ψ in the Hardy-Weinberg mixture model and null (non-HW) model provide quantitative descriptions of the distribution of genotypes and evidence depths in a given population from the perspective of a Hardy-Weinberg equilibrium vs. the alternative non-HW scenario. We can estimate these parameters directly from a set of observed evidence depths using a standard expectation-maximization (EM) algorithm^69^. The resulting estimates of model parameters can be interpreted in ways consistent with concepts from population genetics: allele frequencies (π and α), the positioning of these genotypes in our evidence depth data (μ_2_), the width of these genotypes in our evidence depth data (σ_2_ and σ_3_), and our noise measurement in our null model (λ).

To run the EM algorithm, we first derive maximum likelihood estimators (MLEs) for each model parameter. In the HW model, we use standard MLEs for the Gaussian parameters, noting that the location parameters of the 2nd and 3rd modes are coupled. In the non-HW (null) model, the MLEs are

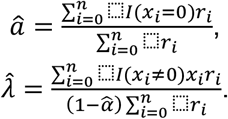

In our analysis, we first determine whether the data fits more closely with the proposed Hardy-Weinberg model or the non-Hardy-Weinberg (null) model using a numerical Kolmogorov-Smirnov (KS) test to evaluate the null hypothesis that the observed data are plausibly generated from a population with *q* = 0. We construct the null distribution of the test statistic numerically in the following manner:

1. Fit the null model to the dataset, *X*, from which we estimate the parameters α^ and λ^.
2. Given these estimated parameters, we compute the empirical distribution function (EDF) for *X* and the cumulative distribution function (CDF), evaluated at each unique value in *X*, using the estimated model parameters α^ and λ^.
3. Calculate the empirical test statistic *D*_∗_ (KS distance) between the EDF and CDF.
4. Then construct the null distribution of *D*. Generate >1000 datasets Ψ_*i*_ from the null model with estimated parameters α^ and λ^; fit the null model to each generated dataset; for each generated dataset, we calculate *D* relative to its own fitted model.
5. Calculate the p-value as the fraction of generated datasets with a test statistic *D* that are at least as large as *D*_∗_.

Staffr can also characterize the degree of overlap of the evidence depths for the relevant genotypes under the HW equilibrium model. This test uses the *common language effect size* or *Mann-Whitney U* statistic (which ranges from 0.5 to 1.0) to quantify the separability of modes in the evidence depth data. The steps of the process are as follows:

1. For each point, assign it to the mode (genotype) with the greatest responsibility *r*_*ij*_.
2. For a large number of rounds, choose two points uniformly at random; let *y* denote the value from the heterozygous mode and let *z* denote the value from the homozygous alternate mode.
3. Let *U* denote the fraction of times that *y* > *z*.

If *U* = 0.5, then modes are statistically indistinguishable, and if *U* = 1.0 then the modes are completely statistically distinguishable, with intermediate values indicating some degree of overlap.

Staffr can also identify outliers relative to a non-HW distribution of evidence, which can be interpreted as evidence for non-reference genotypes. Its method for detecting such outliers is as follows:

1. Over the non-zero evidence values in the data, the MLE of the non-HW model’s geometric distribution parameter λ is given by these data’s mean value.
2. Parameterize the geometric probability mass function and evaluate *f*(*x*|λ) for each unique evidence value *x*.

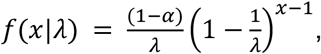
3. Let *b* denote a user-selected minimum number of observations allowed for non-outliers. In practice, *b* = 1 is a relatively inclusive threshold, while *b* = 0.1 is more conservative.
4. Declare all evidence values for which *f*(*x*_⬚_|λ) − *b*/*n* < 0 to be outliers, i.e., values that are statistically unlikely to be observed under the null model.

### Statistics and Visualizations

We conducted all analysis and visualization using python with other packages: Scipy, pandas, numpy, matplotlib, and seaborn. We visualized the SV at read level using the IGV and samplot. Upset package(https://github.com/jnothman/UpSetPlot)was used to visualize the intersection of the mosaic TEs.

## Code Availability

exocrd-lr source code can be found at https://github.com/zhengxinchang/excord-lr. STIX with a long-reads data compatible version can be found at https://github.com/ryanlayer/stix.

## Data Availability

GIAB HG002 PacBio HiFi data(27x) and Oxford Nanopore data(43x) were collected at:https://ftp-trace.ncbi.nlm.nih.gov/ReferenceSamples/giab/data/AshkenazimTrio/HG002_NA24385_son/PacBio_CCS_15kb/ and https://ftp-trace.ncbi.nlm.nih.gov/ReferenceSamples/giab/data/AshkenazimTrio/HG002_NA24385_son/UCSC_Ultralong_OxfordNanopore_Promethion/. GIAB HG002 callset were collected from https://ftp.ncbi.nlm.nih.gov/ReferenceSamples/giab/release/AshkenazimTrio/HG002_NA24385_son/NIST_SV_v0.6/ and https://ftp-trace.ncbi.nlm.nih.gov/ReferenceSamples/giab/data/AshkenazimTrio/analysis/NIST_HG002_DraftBenchmark_defrabbV0.012-20231107/HG00733 callsets with three reference versions are hosted at https://doi.org/10.5281/zenodo.13702318. CMRG bed was downloaded at https://ftp-trace.ncbi.nlm.nih.gov/ReferenceSamples/giab/data/AshkenazimTrio/analysis/NIST_HG002_medical_genes_SV_benchmark_v0.01/. BAM and CRAM files from The 1000 Genomes Project were downloaded from https://s3.amazonaws.com/1000g-ont/index.html?prefix=FIRST_100_FREEZE/minimap2_2.24_alignment_data/, https://s3.amazonaws.com/1000g-ont/index.html?prefix=FIRST_100_FREEZE/minimap2_2.24_chm13_alignment_data/, https://ftp.1000genomes.ebi.ac.uk/vol1/ftp/data_collections/1KG_ONT_VIENNA/hg38/, and https://ftp.1000genomes.ebi.ac.uk/vol1/ftp/data_collections/1KG_ONT_VIENNA/t2t/. BAM files from HPRC project were downloaded from https://human-pangenomics.s3.amazonaws.com/index.html?prefix=working/HPRC/. The COLO829 callset was collected from(https://zenodo.org/records/10819636.). The COSMIC SV were downloaded from https://cancer.sanger.ac.uk/cosmic. The Mosaic TE callset were obtained with request form authors. The raw reads of TP10plus were downloaded from the SRA database(PRJNA636606). Details of the data link can also be found at **Supplementary Table S7**. All software used (with versions) is listed in **Supplementary Table S6.**

## Acknowledgments

We are grateful to the Human Pangenome Reference Consortium, 1000 Genomes Project and the Genome in a Bottle Consortium for releasing datasets essential for this study. The data from the 1000 Genomes ONT Vienna project were generated at the Institute of Molecular Pathology (Vienna, Austria) with funds provided by Boehringer-Ingelheim. Please note that the 1000 Genomes ONT Vienna data, including those generated in this study, are subject to an embargo for genome-wide analyses until the primary publication by Schloissnig et al. (https://www.biorxiv.org/content/10.1101/2024.04.18.590093v1) has completed its peer-review and is published. This embargo is outlined here: https://ftp.1000genomes.ebi.ac.uk/vol1/ftp/data_collections/1KG_ONT_VIENNA/README_1KG_ONT_VIENNA_datareuse_statement_20240227.md. To comply with these restrictions, all genome-wide population data and related datasets from this study have been deposited in the original 1000 Genomes ONT archive at IGSR.

## Contributions

X.Z. developed the STIX LR and performed the analysis. M.C. performed the STIX SR analysis. B.M. & A.C. implemented the mixture model. R.M.L & F.J.S. supervised. All authors contributed to writing and reviewing the manuscript.

## Reference

1. Mahmoud, M. et al. Structural variant calling: the long and the short of it. Genome Biol. 20, 246 (2019).

2. Mahmoud, M. et al. Utility of long-read sequencing for All of Us. Nat. Commun. 15, 1–13 (2024).

3. Sanchis-Juan, A. et al. Complex structural variants in Mendelian disorders: identification and breakpoint resolution using short- and long-read genome sequencing. Genome Med. 10, 95 (2018).

4. Carvalho, C. M. B. & Lupski, J. R. Mechanisms underlying structural variant formation in genomic disorders. Nat. Rev. Genet. 17, 224–238 (2016).

5. Sekar, S. et al. Complex mosaic structural variations in human fetal brains. Genome Res. 30, 1695–1704 (2020).

6. Papaemmanuil, E. et al. Genomic Classification and Prognosis in Acute Myeloid Leukemia. N. Engl. J. Med. 374, 2209–2221 (2016).

7. Cancer Genome Atlas Research Network. Comprehensive genomic characterization of squamous cell lung cancers. Nature 489, 519–525 (2012).

8. Gao, R. et al. Punctuated copy number evolution and clonal stasis in triple-negative breast cancer. Nat. Genet. 48, 1119–1130 (2016).

9. Cosenza, M. R., Rodriguez-Martin, B. & Korbel, J. O. Structural Variation in Cancer: Role, Prevalence, and Mechanisms. Annu. Rev. Genomics Hum. Genet. 23, 123–152 (2022).

10. Dubois, F., Sidiropoulos, N., Weischenfeldt, J. & Beroukhim, R. Publisher Correction: Structural variations in cancer and the 3D genome. Nat. Rev. Cancer (2024) doi:10.1038/s41568-024-00738-y.

11. Jamal-Hanjani, M. et al. Tracking the Evolution of Non-Small-Cell Lung Cancer. N. Engl. J. Med. 376, 2109–2121 (2017).

12. Logsdon, G. A., Vollger, M. R. & Eichler, E. E. Long-read human genome sequencing and its applications. Nat. Rev. Genet. 21, 597–614 (2020).

13. Miller, D. E. et al. Targeted long-read sequencing identifies missing disease-causing variation. Am. J. Hum. Genet. 108, 1436–1449 (2021).

14. Wenger, A. M. et al. Accurate circular consensus long-read sequencing improves variant detection and assembly of a human genome. Nat. Biotechnol. 37, 1155–1162 (2019).

15. Ebert, P. et al. Haplotype-resolved diverse human genomes and integrated analysis of structural variation. Science 372, (2021).

16. McLaren, W. et al. The Ensembl Variant Effect Predictor. Genome Biol. 17, 122 (2016).

17. Wang, K., Li, M. & Hakonarson, H. ANNOVAR: functional annotation of genetic variants from high-throughput sequencing data. Nucleic Acids Res. 38, e164 (2010).

18. Cingolani, P. et al. A program for annotating and predicting the effects of single nucleotide polymorphisms, SnpEff: SNPs in the genome of Drosophila melanogaster strain w1118; iso-2; iso-3. *Fly* 6, 80–92 (2012).

19. Lek, M. et al. Analysis of protein-coding genetic variation in 60,706 humans. Nature 536, 285–291 (2016).

20. Lappalainen, I. et al. DbVar and DGVa: public archives for genomic structural variation. Nucleic Acids Res. 41, D936–41 (2013).

21. Karczewski, K. J. et al. The mutational constraint spectrum quantified from variation in 141,456 humans. Nature 581, 434–443 (2020).

22. Collins, R. L. et al. A structural variation reference for medical and population genetics. Nature 581, 444–451 (2020).

23. Danis, D. et al. SvAnna: efficient and accurate pathogenicity prediction of coding and regulatory structural variants in long-read genome sequencing. Genome Med. 14, 1–13 (2022).

24. Nicholas, T. J., Cormier, M. J. & Quinlan, A. R. Annotation of structural variants with reported allele frequencies and related metrics from multiple datasets using SVAFotate. BMC Bioinformatics 23, 490 (2022).

25. English, A. C., Menon, V. K., Gibbs, R. A., Metcalf, G. A. & Sedlazeck, F. J. Truvari: refined structural variant comparison preserves allelic diversity. Genome Biol. 23, 271 (2022).

26. Chowdhury, M., Pedersen, B. S., Sedlazeck, F. J., Quinlan, A. R. & Layer, R. M. Searching thousands of genomes to classify somatic and novel structural variants using STIX. Nat. Methods 19, 445–448 (2022).

27. Foord, C. et al. The variables on RNA molecules: concert or cacophony? Answers in long-read sequencing. Nat. Methods 20, 20–24 (2023).

28. English, A. C. et al. Analysis and benchmarking of small and large genomic variants across tandem repeats. Nat. Biotechnol. (2024) doi:10.1038/s41587-024-02225-z.

29. Beyter, D. et al. Long-read sequencing of 3,622 Icelanders provides insight into the role of structural variants in human diseases and other traits. Nat. Genet. 53, 779–786 (2021).

30. Gong, J. et al. Long-read sequencing of 945 Han individuals identifies novel structural variants associated with phenotypic diversity and disease susceptibility. medRxiv 2024.03.21.24304654 (2024) doi:10.1101/2024.03.21.24304654.

31. De Coster, W., Weissensteiner, M. H. & Sedlazeck, F. J. Towards population-scale long-read sequencing. Nat. Rev. Genet. 22, 572–587 (2021).

32. Nurk, S. et al. The complete sequence of a human genome. Science 376, 44–53 (2022).

33. Eilbeck, K., Quinlan, A. & Yandell, M. Settling the score: variant prioritization and Mendelian disease. Nat. Rev. Genet. 18, 599–612 (2017).

34. Han, L. et al. Functional annotation of rare structural variation in the human brain. Nat. Commun. 11, 1–13 (2020).

35. Demidov, G. et al. Structural variant calling and clinical interpretation in 6224 unsolved rare disease exomes. Eur. J. Hum. Genet. 1–7 (2024).

36. Layer, R. M. et al. GIGGLE: a search engine for large-scale integrated genome analysis. Nat. Methods 15, 123–126 (2018).

37. Zook, J. M. et al. A robust benchmark for detection of germline large deletions and insertions. Nat. Biotechnol. 38, 1347–1355 (2020).

38. Rang, F. J., Kloosterman, W. P. & de Ridder, J. From squiggle to basepair: computational approaches for improving nanopore sequencing read accuracy. Genome Biol. 19, 90 (2018).

39. Wang, Y., Zhao, Y., Bollas, A., Wang, Y. & Au, K. F. Nanopore sequencing technology, bioinformatics and applications. Nat. Biotechnol. 39, 1348–1365 (2021).

40. Wagner, J. et al. Curated variation benchmarks for challenging medically relevant autosomal genes. Nat. Biotechnol. 40, 672–680 (2022).

41. Smolka, M. et al. Detection of mosaic and population-level structural variants with Sniffles2. Nat. Biotechnol. (2024) doi:10.1038/s41587-023-02024-y.

42. Jiang, T., et al. Regenotyping structural variants through an accurate force-calling method. *bioRxiv* (2022) doi:10.1101/2022.08.29.505534.

43. Fairley, S., Lowy-Gallego, E., Perry, E. & Flicek, P. The International Genome Sample Resource (IGSR) collection of open human genomic variation resources. Nucleic Acids Res. 48, D941–D947 (2020).

44. Liao, W.-W. et al. A draft human pangenome reference. Nature 617, 312–324 (2023).

45. Gustafson, J. A. et al. Nanopore sequencing of 1000 Genomes Project samples to build a comprehensive catalog of human genetic variation. medRxiv (2024) doi:10.1101/2024.03.05.24303792.

46. Wang, T. et al. The Human Pangenome Project: a global resource to map genomic diversity. Nature 604, 437–446 (2022).

47. gnomAD. https://gnomad.broadinstitute.org/help/sv-overview.

48. van Belzen, I. A. E. M., Schönhuth, A., Kemmeren, P. & Hehir-Kwa, J. Y. Structural variant detection in cancer genomes: computational challenges and perspectives for precision oncology. NPJ Precis Oncol 5, 15 (2021).

49. Sun, R., Hu, Z. & Curtis, C. Big Bang Tumor Growth and Clonal Evolution. Cold Spring Harb. Perspect. Med. 8, (2018).

50. Paulin, L. F. et al. The benefit of a complete reference genome for cancer structural variant analysis. medRxiv (2024) doi:10.1101/2024.03.15.24304369.

51. Sondka, Z. et al. COSMIC: a curated database of somatic variants and clinical data for cancer. Nucleic Acids Res. 52, D1210–D1217 (2023).

52. Jaiswal, S. & Ebert, B. L. Clonal hematopoiesis in human aging and disease. Science 366, (2019).

53. Lee-Six, H. et al. The landscape of somatic mutation in normal colorectal epithelial cells. Nature 574, (2019).

54. Pascarella, G. et al. Recombination of repeat elements generates somatic complexity in human genomes. Cell 185, 3025–3040.e6 (2022).

55. Weischenfeldt, J., Symmons, O., Spitz, F. & Korbel, J. O. Phenotypic impact of genomic structural variation: insights from and for human disease. Nat. Rev. Genet. 14, 125–138 (2013).

56. Thibodeau, M. L. et al. Improved structural variant interpretation for hereditary cancer susceptibility using long-read sequencing. Genet. Med. 22, 1892–1897 (2020).

57. Sherry, S. T., Ward, M. & Sirotkin, K. dbSNP-database for single nucleotide polymorphisms and other classes of minor genetic variation. Genome Res. 9, 677–679 (1999).

58. Chen, S. et al. A genomic mutational constraint map using variation in 76,156 human genomes. Nature 625, 92–100 (2024).

59. UK10K Consortium et al. The UK10K project identifies rare variants in health and disease. Nature 526, 82–90 (2015).

60. Cao, Y. et al. The ChinaMAP analytics of deep whole genome sequences in 10,588 individuals. Cell Res. 30, 717–731 (2020).

61. Martincorena, I. et al. Tumor evolution. High burden and pervasive positive selection of somatic mutations in normal human skin. Science 348, 880–886 (2015).

62. Martincorena, I. et al. Somatic mutant clones colonize the human esophagus with age. Science 362, 911–917 (2018).

63. English, A., et al. Benchmarking of small and large variants across tandem repeats. bioRxiv (2023) doi:10.1101/2023.10.29.564632.

64. Byrska-Bishop, M. et al. High-coverage whole-genome sequencing of the expanded 1000 Genomes Project cohort including 602 trios. Cell 185, 3426–3440.e19 (2022).

65. Jeffares, D. C. et al. Transient structural variations have strong effects on quantitative traits and reproductive isolation in fission yeast. Nat. Commun. 8, 14061 (2017).

66. Pedersen, B. S. & Quinlan, A. R. Mosdepth: quick coverage calculation for genomes and exomes. Bioinformatics 34, 867–868 (2018).

67. Li, H. Minimap2: pairwise alignment for nucleotide sequences. Bioinformatics 34, 3094– 3100 (2018).

68. Feng, C. X. A comparison of zero-inflated and hurdle models for modeling zero-inflated count data. J Stat Distrib Appl 8, 8 (2021).

69. Moon, T. K. The expectation-maximization algorithm. IEEE Signal Process. Mag. 13, 47–60 (1996).

70. Olson, N. D. precisionFDA Truth Challenge V2: Calling variants from short- and long-reads in difficult-to-map regions. National Institute of Standards and Technology 10.18434/MDS2-2336 (2020)

